## Supplementary Table S12 for "Positive selection and relaxed purifying selection contribute to rapid evolution of male-biased genes in a dioecious flowering plant"

**Table S12** Functions and References associated with abiotic stress and immune responses, organ developments of male-biased genes under significant positive selection (*P-value* < 0.05) in floral buds.

| OGs | Function Description | Signal* | Stress responses/Developments | References |
|---|---|---|---|---|
| **OG0007149** | mitogen-activated protein kinase kinase kinase 18 (MAPKKK18) | A, B, C, R | drought, heavy metal, microtubule stabilization | Mondal (2022); Zhang and Zhang (2022); Xu and Zhang (2015); Lin, L *et al.* (2021) |
| **OG0007167** | ubiquitin-conjugating enzyme E2 24 (UBC24) | A, B, C, R | pathogens, limited nutrient, heavy metal, male fertility | Unver *et al.* (2013); Sharma *et al.* (2021); Wang *et al.* (2020) |
| OG0007172 | homeobox-DDT domain protein RLT2 (RLT2) | A, B, C, R | plant vegetative phase | Li *et al.* (2012) |
| ***OG0007198*** | la-related protein 6A (LARP6A) | A, B, C, R | salt, mature pollen grain, male gametogenesis | De Vos *et al.* (2019); Bousquet-Antonelli (2021); Zhang *et al.* (2017) |
| OG0009257 | uncharacterized protein | A, B, C, R | | |
| **OG0008422** | regulator of nonsense transcripts UPF2 (UPF2) | A, B, C, R | pathogens, normal growth | Shi *et al.* (2012) |
| **OG0006902** | heat stress transcription factor B-3 (HSFB3) | A, B, C, R | heat, drought, dehydration, pathogens | Mehta *et al.* (2021); Liao *et al.* (2022); Wang *et al.* (2018); Scharf *et al.* (2012) |
| OG0007054 | uncharacterized protein | A, B, C, R | | |
| OG0007066 | suppressor of RPS4-RLD 1 (SRFR1) | A, B, C, R | pathogens | Kwon *et al.* (2004); Kwon *et al.* (2009) |
| OG0007085 | LMBR1 domain-containing protein (LMBR1) | A, B, C, R | | |
| **OG0008757** | phytochrome B (PHYB) | A, B, C, R | heat, cold, drought, red light, jasmonic acid (JA), salicylic acid (SA), root, stem, leaf, flower | Kim *et al.* (2021); Ikeda *et al.* (2021) |
| **OG0007324** | autophagy-related protein 18a (ATG18a) | A, B, C, R | pathogens, nutrient starvation, dark, cold, drought, salt, heavy metal, oxidation, osmotic, jasmonic acid (JA), salicylic | Qi *et al.* (2021); Zhou *et al.* (2015); Su *et al.* (2020); Avin- |

| | | | | |
|---|---|---|---|---|
| | | | acid (SA), senescence | Wittenberg (2019); Xiong *et al.* (2005) |
| OG0009150 | MND1-interacting protein 1 (MIP1) | A, B, C, R | | |
| ***OG0009434*** | carbohydrate esterase (At3g16150) | A, B, C, R | pollen development | Honys & Twell, (2003) |
| **OG0009468** | nucleoredoxin 2 | A, B, C, R | drought, oxidative | Kneeshaw *et al.* (2017); Kotrade *et al.* (2019) |
| **OG0007918** | aminoacylase-1 (ACY1) | A, B, C, R | pathogens, salt, drought, jasmonic acid (JA), salicylic acid (SA), plant growth | Chen *et al.* (2021) |
| **OG0007693** | pheophorbide a oxygenase (PAO) | A, B, C, R | wounding, cold, salt, pathogens, seed, suppression of cell death | Ma *et al.* (2012); Chung *et al.* (2006) |
| **OG0008053** | CHROMATIN REMODELING 5 (CHR5) | A, B, C, R | heat, salt, drought, ABA, pathogens | Song *et al.* (2021) |
| **OG0006505** | E3 ubiquitin-protein ligase RHB1A (RHB1A) | A, B, C, R | pathogens | Shahinnia *et al.* (2022) |
| **OG0006517** | protein STAY-GREEN (SGR) | A, B, C, R | drought, heat | Abdelrahman *et al.* (2017); Faye *et al.* (2022) |
| **OG0008678** | serine carboxypeptidase (SCP) | A, B, C, R | pathogens, light, cold, wound, salt, drought | Liu *et al.* (2022); Xu *et al.* (2021) |
| OG0008965 | sphingolipid transporter spinster homolog 2 (SPNS2) | A, B, C, R | | |
| ***OG0009362*** | 14-3-3 protein 7-like | A, B, C, R | cold, osmotic, nutrient starvation, starch accumulation | Xu *et al.* (2012); Roberts *et al.* (2002); Datta *et al.* (2001) |
| ***OG0009379*** | plant homeodomain (PHD) finger protein 3 (MALE STERILITY 3) | A, B, C, R | salt, ABA hormonal, humidity, light, pollen, pollen grain, tapetum | Wang, QQ *et al.* (2015); Hou *et al.* (2022); Ito and Shinozaki (2002); Ito *et al.* (2007); Li *et al.* (2011) |
| **OG0006703** | nudix hydrolase 8-like protein (NUDT8) | A, B, C, R | pathogens | Fonseca and Dong (2014) |
| **OG0008506** | Nicotinamide adenine dinucleotide kinase 2 (NADK2) | A, B, C, R | heavy metal, salt, heat, cold, light, drought, hormonal, oxidative, pathogens, root, stem, leaf | Li *et al.* (2018); Hashida *et al.* (2009); Hashida *et al.* (2007); Gakière *et al.* |

| | | | | |
|---|---|---|---|---|
| | | | | (2018); Takahara *et al.* (2010); Chai *et al.* (2005); Takahashi *et al.* (2006); Pétriacq *et al.* (2013) |
| **OG0008891** | homeobox-leucine zipper protein ATHB-12 (ATHB-12) | A, B, C | water, salt | Lee and Chun (1998); Shin *et al.* (2004) |
| **OG0006931** | copper-transporting ATPase HMA5 (HMA5) | A, B, C | heavy metal | Ma *et al.* (2022) |
| **OG0009171** | STRICTOSIDINE SYNTHASE-LIKE 4 (SSL4) | A, B, C | heat, pathogens | Sohani *et al.* (2009); Aoun *et al.* (2020) |
| **OG0007942** | isocitrate lyase | A, B, C | salt | Yuenyong *et al.* (2019) |
| OG0009067 | Acyl-coenzyme A oxidase, peroxisomal | A, B, C | | |
| **OG0008247** | Homogentisate phytyltransferase 1 (HPT1) | A, B, C | light, oxidative, nutrient starvation | Collakova and DellaPenna (2003) |
| **OG0007434** | BTB/POZ domain-containing protein (BTB/POZ) | A, B, R | pathogens | Zhao *et al.* (2022) |
| **OG0009245** | zinc transporter 11 (ZIP11) | A, B, R | heavy metal | Milner *et al.* (2013) |
| **OG0006058** | phosphatase 2C 16 (HAB1) | A, B, R | abscisic acid (ABA), cold, drought, osmotic, salinity, bacteria, fungi | Singh *et al.* (2016) |
| **OG0006604** | DEHYDRATION-INDUCED 19 homolog 3 (DI19-3) | A, B, R | drought, salinity | (Milner *et al.*, 2013) |
| OG0006674 | SH3 domain-containing YSC84-like protein | A, B, R | | |
| OG0008723 | uncharacterized protein | A, B, R | | |
| **OG0009128** | phospholipase D beta 1 (PLDBETA1) | A, B, R | salt, abscisic acid (ABA), cold, dehydration, drought | Hong *et al.* (2016); Laxalt *et al.* (2001) |
| **OG0009144** | beta-amylase 3 (BAM3) | A, B, R | cold | Sun *et al.* (2021); |
| OG0008799 | 1,2-dihydroxy-3-keto-5-methylthiopentene dioxygenase 2 | A, B, R | | |
| OG0006256 | UPF0496 protein | A, B, R | | |
| **OG0007963** | Inositol pentakisphosphate 2 kinase (ITPK1) | A, B, R | heavy metal, nutrient starvation, salicylic acid (SA), pathogens | Sun *et al.* (2016) |
| OG0008345 | uncharacterized protein | A, B, R | | |

| | | | | |
|---|---|---|---|---|
| OG0006863 | cleft lip and palate transmembrane protein | A, B, R | | |
| **OG0009312** | 4-hydroxyphenylpyruvate dioxygenase (HPD) | A, B, R | oxidative | Falk *et al.* (2002) |
| **OG0007860** | BTB/POZ domain-containing protein (BTB/POZ) | A, B, R | pathogens | Zhao *et al.* (2022) |
| OG0009654 | 1-phosphatidylinositol-3-phosphate 5-kinase FAB1C (FAB1C) | A, B, R | | |
| OG0007874 | Lysine-specific demethylase rbr-2 | A, B, R | | |
| **OG0008241** | calcium-transporting ATPase 8 (ACA8) | A, B, R | salt | Gong *et al.* (2005) |
| OG0008522 | flocculation protein FLO11 | A, B, R | | |
| **OG0007822** | trihelix transcription factor GT-2 (GT-2) | A, C, R | salt, hormonal, pathogen, trichome | Zhu *et al.* (2022); Breuer *et al.* (2009); Kaplan-Levy *et al.* (2012); Volz *et al.* (2018) |
| **OG0008751** | chloroplast unusual positioning protein 1 (CHUP1) | B, C, R | light | Morita and Nakamura (2012) |
| **OG0007594** | UPF0051 protein ABCI8 (ABCI8) | B, C, R | light, salt | Capriotti *et al.* (2014) |
| **OG0009517** | LRR receptor-like serine threonine protein kinase（LRR-RLK) | A, B | pathogen, hormonal, drought, salt, cold, root, floral organ, ovule, anther | Afzal *et al.* (2008); Morris and Walker (2003); Ye *et al.* (2017); Hosseini *et al.* (2020) |
| OG0009539 | hypothetical protein | A, B | | |
| ***OG0006312*** | Glutaredoxin-C6 | A, B | anther development, male gametogenesis, | Xing and Zachgo (2008); Hong *et al.* (2012); Stroher and Millar (2012) |
| OG0008196 | protein MEI2-like 2 | A, B | | |
| OG0008439 | transcription factor DIVARICATA | A, B | | |
| **OG0006928** | cationic amino acid transporter 2, vacuolar | A, B | cold, salt, stamen | Feng *et al.* (2018); Yang *et al.* (2013) |
| **OG0007317** | E3 ubiquitin-protein ligase RHB1A | A, B | pathogens | Shahinnia *et al.* (2022) |
| **OG0009191** | SCY1-like protein 2 | A, B | pathogens | Yao *et al.* (2022); |
| **OG0008092** | $K^+$ efflux antiporter 3 | A, B | drought, salt, alkaline | Sheng *et al.* (2014); Saddhe *et* |

| | | | | |
|---|---|---|---|---|
| | | | | *al.* (2021); Zhu *et al.* (2018) |
| OG0008351 | MLP1, MLP2-like protein | A, B | | |
| OG0008659 | monofunctional riboflavin biosynthesis RIBA3 (RIBA3) | A, B | | |
| **OG0007568** | quinolinate synthase (QS) | A, B | salt | Wei *et al.* (2020) |
| **OG0006141** | signal peptide peptidase-like 4 (SPPL4) | A, B | pathogens | Pinter *et al.* (2019) |
| OG0007820 | uncharacterized protein | A, B | | |
| ***OG0008526*** | Plant U-box domain-containing protein 4 (PUB4) | A, B | drought, tapetal cells, pollen development, starch deposition | Adler *et al.* (2017); Wang *et al.* (2013); Trenner *et al.* (2022); Zhang, WD *et al.* (2020) |
| OG0008469 | uncharacterized protein | A, C | | |
| OG0009423 | protein phosphatase 2C 72 | A, C | | |
| OG0006198 | uncharacterized protein | A, C | | |
| OG0006967 | mitoferrin-like protein | A, R | | |
| OG0006239 | uncharacterized protein | A, R | | |
| OG0009068 | uncharacterized protein | A, R | | |
| OG0008403 | WAT1-related protein | A | | |
| ***OG0008456*** | Glutaredoxin | A | heat, oxidative, anther, pollen | Xing and Zachgo (2008); Stroher and Millar (2012); Hong *et al.* (2012) |
| **OG0007047** | lysM domain receptor-like kinase 4 (LYK4) | A | fungal, salt | Wan *et al.* (2008); Buendia *et al.* (2018); Wan *et al.* (2012) |
| **OG0008731** | WRKY transcription factor 45 (WRKY45) | A | drought, salt, heat, ABA, nutrient starvation, pathogens | Chen *et al.* (2017); Wang *et al.* (2014); Chen *et al.* (2012); Li, WX *et al.* (2020) |
| **OG0007380** | RPM1-interacting protein 4 (RIN4) | A | pathogens | Prokchorchik *et al.* (2020) |
| OG0006802 | plant intracellular Ras-group-related LRR protein 6 (PIRL6) | A | | |
| **OG0006130** | LOB domain-containing protein 19 (LBD19) | A | heat, light, pathogens, callus formation | Zhang, YW *et al.* (2020); Liu *et al.* (2019) |
| ***OG0009273*** | autophagy-related protein 18f | A | pathogens, nutrient starvation, | Qi *et al.* (2021); |

| | | | | |
|---|---|---|---|---|
| | (ATG18f) | | drought, heat, cold, salt, heavy metal, oxidative, anther, pollen, root, stem, leaf | Li, SH *et al.* (2020); Avin-Wittenberg (2019); Zhou *et al.* (2015); Wang, Y *et al.* (2015) |
| ***OG0007429*** | protein JINGUBANG-like | A | jasmonic acid (JA), pollen grain | Ju *et al.* (2016) |
| OG0009210 | leucine-rich repeat receptor-like tyrosine-protein kinase PXC3 (PXC3) | A | | |
| **OG0006776** | ASPARTIC PROTEASE IN GUARD CELL 1 (ASPG1) | A | drought, pathogens | Yao *et al.* (2012); Sebastian *et al.* (2020); Huang *et al.* (2013); Gao *et al.* (2017); Figueiredo *et al.* (2021) |
| ***OG0008560*** | CLAVATA3 (CLV3) /ESR (CLE)-related protein 13 (CLV3/CLE13) | B | cold, heat, drought, salt, cytokinin, hormonal, floral organ, stamen, anther, root, shoot | Zhang *et al.* (2022); Lin, H *et al.* (2021); Cui *et al.* (2022); Yamaguchi *et al.* (2016); Jun *et al.* (2010); Laffont *et al.* (2020) |
| **OG0008786** | basic 7S globulin-like protein | C | antibacterial, allergen, heat | Hirano (2021) |
| **OG0006303** | zinc finger CCCH domain-containing protein 20 (C3H20/TZF2) | R | salt, drought, flooding, cold, oxidative, ABA, JA, flower | Han *et al.* (2021); Lee *et al.* (2012); Bogamuwa and Jang (2014); Huang *et al.* (2012); Huang *et al.* (2011) |
| OG0006084 | uncharacterized protein | R | | |
| OG0008460 | Pectinesterase inhibitor 45 (PME45) | R | | |
| **OG0006683** | auxin-binding protein ABP19a (ABP19a) | R | dehydration, drought | Vessal *et al.* (2020) |
| **OG0007044** | ABC transporter F family member 5 (ABCF5) | R | dry | Hwang *et al.* (2016) |
| OG0006937 | uncharacterized protein | R | | |
| **OG0009424** | zinc finger CCCH domain-containing protein 20 (C3H20/TZF2) | R | salt, drought, flooding, cold, oxidative, ABA, JA, flower | Bogamuwa and Jang (2014); Lee *et al.* (2012); Han *et al.* (2021); Huang *et al.* |

| | | | | |
|---|---|---|---|---|
| | | | | (2012); Huang *et al.* (2011) |
| **OG0008325** | E3 ubiquitin-protein ligase RING1 | R | salt, drought, cold, heat, bacterial | Wang *et al.* (2022); Marino *et al.* (2012) |
| **OG0009024** | BTB/POZ domain and ankyrin repeat-containing protein NPR2 | R | pathogens | Pajerowska-Mukhtar *et al.* (2013) |
| **OG0006402** | nucleoredoxin 1 | R | salt, drought | Zhang *et al.* (2014); Zhang *et al.* (2021) |

OGs in bold font are associated with biotic and abiotic stress responses. OGs in bold and italic fonts are associated with biotic and abiotic stress responses, pollen developments and maturation.

Asterisk (*) indicates that positive selections were tested by aBSREL (A), BUSTED (B), CodeML (C) and RELAX (R).


**Abdelrahman M, El-Sayed M, Jogaiah S, Burritt DJ, Tran LSP. 2017.** The "STAY-GREEN" trait and phytohormone signaling networks in plants under heat stress. *Plant Cell Reports* **36**: 1009-1025.

**Adler G, Konrad Z, Zamir L, Mishra AK, Raveh D, Bar-Zvi D. 2017.** The *Arabidopsis* paralogs, *PUB46* and *PUB48*, encoding U-box E3 ubiquitin ligases, are essential for plant response to drought stress. *BMC Plant Biology* **17**: 8.

**Afzal AJ, Wood AJ, Lightfoot DA. 2008.** Plant receptor-like serine threonine kinases: roles in signaling and plant defense. *Molecular Plant-Microbe Interactions* **21**: 507-517.

**Aoun N, Desaint H, Boyrie L, Bonhomme M, Deslandes L, Berthomé R, Roux F. 2020.** A complex network of additive and epistatic quantitative trait loci underlies natural variation of *Arabidopsis thaliana* quantitative disease resistance to *Ralstonia solanacearum* under heat stress. *Molecular Plant Pathology* **21**: 1405-1420.

**Avin-Wittenberg T. 2019.** Autophagy and its role in plant abiotic stress management. *Plant Cell and Environment* **42**: 1045-1053.

**Bogamuwa SP, Jang JC. 2014.** Tandem CCCH Zinc finger proteins in plant growth, development and stress response. *Plant and Cell Physiology* **55**: 1367-1375.

**Bousquet-Antonelli C. 2021.** LARP6 proteins in plants. *Biochemical Society Transactions* **49**: 1975-1983.

**Breuer C, Kawamura A, Ichikawa T, Tominaga-Wada R, Wada T, Kondou Y, Muto S, Matsui M, Sugimoto K. 2009.** The trihelix transcription factor GTL1 regulates ploidy-dependent cell growth in the *Arabidopsis* trichome. *Plant Cell* **21**: 2307-2322.

**Buendia L, Girardin A, Wang TM, Cottret L, Lefebvre B. 2018.** LysM receptor-like kinase and LysM receptor-Like protein families: an update on phylogeny and functional characterization. *Frontiers in Plant Science* **9**: 1531.

**Capriotti AL, Borrelli GM, Colapicchioni V, Papa R, Piovesana S, Samperi R, Stampachiacchiere S, Lagana A. 2014.** Proteomic study of a tolerant genotype of durum wheat under salt-stress conditions. *Analytical and Bioanalytical Chemistry* **406**: 1423-1435.

**Chai MF, Chen QJ, An R, Chen YM, Chen J, Wang XC. 2005.** NADK2, an *Arabidopsis* chloroplastic NAD kinase, plays a vital role in both chlorophyll synthesis and chloroplast protection. *Plant Molecular Biology* **59**: 553-564.

**Chen DB, Li JH, Jiao FC, Wang QQ, Li J, Pei YH, Zhao M, Song XY, Guo XM. 2021.** *ZmACY-1* antagonistically regulates growth and stress responses in *Nicotiana benthamiana*. *Frontiers in Plant Science* **1212**: 593001.

**Chen F, Hu Y, Vannozzi A, Wu KC, Cai HY, Qin Y, Mullis A, Lin ZG, Zhang LS. 2017.** The WRKY transcription factor family in model plants and crops. *Critical Reviews in Plant Sciences* **36**: 311-335.

**Chen LG, Song Y, Li SJ, Zhang LP, Zou CS, Yu DQ. 2012.** The role of WRKY transcription factors in plant abiotic stresses. *Biochimica Et Biophysica Acta-Gene Regulatory Mechanisms* **1819**: 120-128.

**Chung DW, Pruzinska A, Hortensteiner S, Ort DR. 2006.** The role of pheophorbide a oxygenase expression and activity in the canola green seed problem. *Plant Physiology* **142**: 88-97.

**Collakova E, DellaPenna D. 2003.** The role of homogentisate phytyltransferase and other tocopherol pathway enzymes in the regulation of tocopherol synthesis during abiotic stress. *Plant Physiology* **133**: 930-940.

**Cui YW, Lu XT, Gou XP. 2022.** Receptor-like protein kinases in plant reproduction: current understanding and future perspectives. *Plant Communications* **3:** 100273.

**Datta R, Chourey PS, Pring DR, Tang HV. 2001.** Gene-expression analysis of sucrose-starch metabolism during pollen maturation in cytoplasmic male-sterile and fertile lines of sorghum. *Sexual Plant Reproduction* **14**: 127-134.

**De Vos S, Van Stappen G, Sorgeloos P, Vuylsteke M, Rombauts S, Bossier P. 2019.** Identification of salt stress response genes using the *Artemia* transcriptome. *Aquaculture* **500**: 305-314.

**Falk J, Krauss N, Dahnhardt D, Krupinska K. 2002.** The senescence associated gene of barley encoding 4-hydroxyphenylpyruvate dioxygenase is expressed during oxidative stress. *Journal of Plant Physiology* **159**: 1245-1253.

**Faye JM, Akata EA, Sine B, Diatta C, Cisse N, Fonceka D, Morris GP. 2022.** Quantitative and population genomics suggest a broad role of stay-green loci in the drought adaptation of sorghum. *Plant Genome* **15:** e20176.

**Feng L, Yang TY, Zhang ZL, Li FD, Chen Q, Sun J, Shi CY, Deng WW, Tao MM, Tai YL, Yang H, Cao Q, Wan XC. 2018.** Identification and characterization of cationic amino acid transporters (CATs) in tea plant (*Camellia sinensis*). *Plant Growth Regulation* **84**: 57-69.

**Figueiredo L, Santos RB, Figueiredo A. 2021.** Defense and offense otrategies: the role of aspartic proteases in plant-pathogen interactions. *Biology* **10:** 75.

**Fonseca JP, Dong XN. 2014.** Functional characterization of a nudix hydrolase *AtNUDX8* upon pathogen attack indicates a positive role in plant immune responses. *PLoS One* **9:** e114119.

**Gakière B, Hao J, de Bont L, Pétriacq P, Nunes-Nesi A, Fernie AR. 2018.** NAD+ biosynthesis and signaling in plants. *Critical Reviews in Plant Sciences* **37**: 259-307.

**Gao H, Zhang YH, Wang WL, Zhao KK, Liu CM, Bai L, Li R, Guo Y. 2017.** Two membrane-anchored aspartic proteases contribute to pollen and ovule development. *Plant Physiology* **173**: 219-239.

**Gong QQ, Li PH, Ma SS, Rupassara SI, Bohnert HJ. 2005.** Salinity stress adaptation competence in the extremophile *Thellungiella halophila* in comparison with its relative *Arabidopsis thaliana*. *Plant Journal* **44**: 826-839.

**Han GL, Qiao ZQ, Li YX, Wang CF, Wang BS. 2021.** The roles of CCCH zinc-finger proteins in plant abiotic stress tolerance. *International Journal of Molecular Sciences* **22**: 8327.

**Hashida SN, Takahashi H, Uchimiya H. 2009.** The role of NAD biosynthesis in plant development and stress responses. *Annals of Botany* **103**: 819-824.

**Hirano H. 2021.** Basic 7S globulin in plants. *Journal of Proteomics* **240**: 104209.

**Hong LL, Tang D, Zhu KM, Wang KJ, Li M, Cheng ZK. 2012.** Somatic and reproductive cell development in rice anther is regulated by a putative glutaredoxin. *Plant Cell* **24**: 577-588.

**Hong YY, Zhao J, Guo L, Kim SC, Deng XJ, Wang GL, Zhang GY, Li MY, Wang XM. 2016.** Plant phospholipases D and C and their diverse functions in stress responses. *Progress in Lipid Research* **62**: 55-74.

**Honys D, Twell D. 2003.** Comparative analysis of the *Arabidopsis* pollen transcriptome. *Plant Physiology* **132**: 640-652.

**Hosseini S, Schmidt EDL, Bakker FT. 2020.** Leucine-rich repeat receptor-like kinase II phylogenetics reveals five main clades throughout the plant kingdom. *Plant Journal* **103**: 547-560.

**Hou JJ, Fan WW, Ma RR, Li B, Yuan ZH, Huang WX, Wu YY, Hu Q, Lin CJ, Zhao XQ, Peng B, Zhao LM, Zhang CB, Sun LJ. 2022.** *MALE STERILITY 3* encodes a plant homeodomain-finger protein for male fertility in soybean. *Journal of Integrative Plant Biology* **64**: 1076-1086.

**Huang JY, Zhao XB, Cheng K, Jiang YH, Ouyang YD, Xu CG, Li XH, Xiao JH, Zhang QF. 2013.** OsAP65, a rice aspartic protease, is essential for male fertility and plays a role in pollen germination and pollen tube growth. *Journal of Experimental Botany* **64**: 3351-3360.

**Huang P, Chung MS, Ju HW, Na HS, Lee DJ, Cheong HS, Kim CS. 2011.** Physiological characterization of the *Arabidopsis thaliana* oxidation-related zinc finger 1, a plasma membrane protein involved in oxidative stress. *Journal of plant research* **124**: 699-705.

**Huang P, Ju HW, Min JH, Zhang X, Chung JS, Cheong HS, Kim CS. 2012.** Molecular and physiological characterization of the *Arabidopsis thaliana* oxidation-related zinc finger 2, a plasma membrane protein involved in ABA and salt stress response through the ABI2-mediated signaling pathway. *Plant and Cell Physiology* **53**: 193-203.

**Hwang JU, Song WY, Hong D, Ko D, Yamaoka Y, Jang S, Yim S, Lee E, Khare D, Kim K, Palmgren M, Yoon HS, Martinoia E, Lee Y. 2016.** Plant ABC transporters enable many unique aspects of a terrestrial plants lifestyle. *Mololecular Plant* **9**: 338-355.

**Ikeda H, Suzuki T, Oka Y, Gustafsson ALS, Brochmann C, Mochizuki N, Nagatani A. 2021.** Divergence in red light responses associated with thermal reversion of phytochrome B between high- and low-latitude species. *New Phytologist* **231**: 75-84.

**Ito T, Nagata N, Yoshiba Y, Ohme-Takagi M, Ma H, Shinozaki K. 2007.** Arabidopsis *MALE STERILITY1* encodes a PHD-type transcription factor and regulates pollen and tapetum development. *Plant Cell* **19**: 3549-3562.

**Ito T, Shinozaki K. 2002.** The *MALE STERILITY1* gene of *Arabidopsis*, encoding a nuclear protein with a PHD-finger motif, is expressed in tapetal cells and is required for pollen maturation. *Plant and*

*Cell Physiology* **43**: 1285-1292.

**Ju Y, Guo L, Cai Q, Ma F, Zhu QY, Zhang Q, Sodmergen. 2016.** Arabidopsis JINGUBANG is a negative regulator of pollen germination that prevents pollination in moist environments. *Plant Cell* **28**: 2131-2146.

**Jun J, Fiume E, Roeder AHK, Meng L, Sharma VK, Osmont KS, Baker C, Ha CM, Meyerowitz EM, Feldman LJ, Fletcher JC. 2010.** Comprehensive analysis of CLE polypeptide signaling gene expression and overexpression activity in *Arabidopsis*. *Plant Physiology* **154**: 1721-1736.

**Jung JY, Lee DW, Ryu SB, Hwang I, Schachtman DP. 2017.** *SCYL2* genes are involved in Clathrin-mediated vesicle trafficking and essential for plant growth. *Plant Physiology* **175**: 194-209.

**Kaplan-Levy RN, Brewer PB, Quon T, Smyth DR. 2012.** The trihelix family of transcription factors - light, stress and development. *Trends in Plant Science* **17**: 163-171.

**Kim JY, Lee JH, Park CM. 2021.** A multifaceted action of phytochrome B in plant environmental adaptation. *Frontiers in Plant Science* **12**.

**Kneeshaw S, Keyani R, Delorme-Hinoux V, Imrie L, Loake GJ, Le Bihan T, Reichheld JP, Spoel SH. 2017.** Nucleoredoxin guards against oxidative stress by protecting antioxidant enzymes. *Proceedings of the National Academy of Sciences of the United States of America* **114**: 8414-8419.

**Kotrade P, Sehr EM, Brüggemann W. 2019.** Expression profiles of 12 drought responsive genes in two European (deciduous) oak species during a two-year drought experiment with consecutive drought periods. *Plant Gene* **19**: 100193.

**Kwon SI, Kim SH, Bhattacharjee S, Noh JJ, Gassmann W. 2009.** *SRFR1*, a suppressor of effector-triggered immunity, encodes a conserved tetratricopeptide repeat protein with similarity to transcriptional repressors. *Plant Journal* **57**: 109-119.

**Kwon SI, Koczan JM, Gassmann W. 2004.** Two *Arabidopsis srfr* (suppressor of *rps4-RLD*) mutants exhibit *avrRps4*-specific disease resistance independent of *RPS4*. *Plant Journal* **40**: 366-375.

**Laffont C, Ivanovici A, Gautrat P, Brault M, Djordjevic MA, Frugier F. 2020.** The NIN transcription factor coordinates CEP and CLE signaling peptides that regulate nodulation antagonistically. *Nature Communication* **11**: 3167.

**Laxalt AM, ter Riet B, Verdonk JC, Parigi L, Tameling WIL, Vossen J, Haring M, Musgrave A, Munnik T. 2001.** Characterization of five tomato phospholipase D cDNAs: rapid and specific expression of LePLD beta 1 on elicitation with xylanase. *Plant Journal* **26**: 237-247.

**Lee SJ, Jung HJ, Kang H, Kim SY. 2012.** Arabidopsis zinc finger proteins AtC3H49/AtTZF3 and AtC3H20/AtTZF2 are involved in ABA and JA responses. *Plant and Cell Physiology* **53**: 673-686.

**Lee YH, Chun JY. 1998.** A new homeodomain-leucine zipper gene from *Arabidopsis thaliana* induced by water stress and abscisic acid treatment. *Plant Molecular Biology* **37**: 377-384.

**Li BB, Wang X, Tai L, Ma TT, Shalmani A, Liu WT, Li WQ, Chen KM. 2018.** NAD Kinases: metabolic targets controlling redox co-enzymes and reducing power partitioning in plant stress and development. *Frontiers in Plant Science* **9**: 379.

**Li G, Zhang JW, Li JQ, Yang ZN, Huang H, Xu L. 2012.** Imitation switch chromatin remodeling factors and their interacting RINGLET proteins act together in controlling the plant vegetative phase in *Arabidopsis*. *Plant Journal* **72**: 261-270.

**Li H, Yuan Z, Vizcay-Barrena G, Yang CY, Liang WQ, Zong J, Wilson ZA, Zhang DB. 2011.** *PERSISTENT TAPETAL CELL1* encodes a PHD-finger protein that is required for tapetal cell death and pollen development in rice. *Plant Physiology* **156**: 615-630.

**Li SH, Yan H, Mei WM, Tse YC, Wang H. 2020.** Boosting autophagy in sexual reproduction: a plant perspective. *New Phytologist* **226**: 679-689.

**Li WX, Pang SY, Lu ZG, Jin B. 2020.** Function and mechanism of WRKY transcription factors in abiotic stress responses of plants. *Plants* **9**: 1515.

**Liao Y, Liu Z, Gichira AW, Yang M, Mbichi RW, Meng L, Wan T. 2022.** Deep evaluation of the evolutionary history of the Heat Shock Factor (HSF) gene family and its expansion pattern in seed plants. *PeerJ* **10**: e13603.

**Lin H, Wang W, Chen XG, Sun ZT, Han XL, Wang S, Li Y, Ye WW, Yin ZJ. 2021.** Molecular traits and functional analysis of the CLAVATA3/Endosperm surrounding region-related small signaling peptides in three species of *Gossypium* Genus. *Frontiers in Plant Science* **12**: 671626.

**Lin L, Wu J, Jiang MY, Wang YP. 2021.** Plant mitogen-activated protein kinase cascades in environmental stresses. *International Journal of Molecular Sciences* **22**: 1543.

**Liu SQ, Wang B, Li XJ, Pan JX, Qian XX, Yu YH, Xu P, Zhu J, Xu XF. 2019.** Lateral Organ Boundaries Domain 19 (LBD19) negative regulate callus formation in *Arabidopsis*. *Plant Cell Tissue and Organ Culture* **137**: 485-494.

**Liu YL, Ce F, Tang H, Tian GF, Yang L, Qian W, Dong HL. 2022.** Genome-wide analysis of the serine carboxypeptidase-like (SCPL) proteins in *Brassica napus* L. *Plant Physiology and Biochemistry*

**186**: 310-321.

**Ma N, Ma X, Li AF, Cao XC, Kong LR. 2012.** Cloning and expression analysis of Wheat pheophorbide a oxygenase gene *TaPaO*. *Plant Molecular Biology Reporter* **30**: 1237-1245.

**Ma Y, Liu KC, Zhang CY, Lin F, Hu WB, Jiang Y, Tao XL, Han YL, Han LT, Liu C. 2022.** Comparative root transcriptome analysis of two soybean cultivars with different cadmium sensitivities reveals the underlying tolerance mechanisms. *Genome* **65**: 27-42.

**Marino D, Peeters N, Rivas S. 2012.** Ubiquitination during plant immune signaling. *Plant Physiology* **160**: 15-27.

**Mehta S, Chakraborty A, Roy A, Singh IK, Singh A. 2021.** Fight hard or die trying: current status of lipid signaling during plant-pathogen interaction. *Plants* **10**: 1098.

**Milner MJ, Seamon J, Craft E, Kochian LV. 2013.** Transport properties of members of the ZIP family in plants and their role in Zn and Mn homeostasis. *Journal of Experimental Botany* **64**: 369-381.

**Mondal S. 2022.** Heavy metal stress-induced activation of mitogen-activated protein kinase signalling cascade in plants. *Plant Molecular Biology Reporter* https://doi.org/10.1007/s11105-022-01350-w.

**Morita MT, Nakamura M. 2012.** Dynamic behavior of plastids related to environmental response. *Current Opinion in Plant Biology* **15**: 722-728.

**Morris ER, Walker JC. 2003.** Receptor-like protein kinases: the keys to response. *Current Opinion in Plant Biology* **6**: 339-342.

**Pétriacq P, de Bont L, Tcherkez G, Gakière B. 2013.** NAD: not just a pawn on the board of plant-pathogen interactions. *Plant Signaling and Behavior* **8**: e22477.

**Pajerowska-Mukhtar KM, Emerine DK, Mukhtar MS. 2013.** Tell me more: roles of NPRs in plant immunity. *Trends in Plant Science* **18**: 402-411.

**Pinter N, Hach CA, Hampel M, Rekhter D, Zienkiewicz K, Feussner I, Poehlein A, Daniel R, Finkernagel F, Heimel K. 2019.** Signal peptide peptidase activity connects the unfolded protein response to plant defense suppression by *Ustilago maydis*. *Plos Pathogens* **15:** e1007734.

**Prokchorchik M, Choi S, Chung EH, Won K, Dangl JL, Sohn KH. 2020.** A host target of a bacterial cysteine protease virulence effector plays a key role in convergent evolution of plant innate immune system receptors. *New Phytologist* **225**: 1327-1342.

**Qi H, Xia FN, Xiao S. 2021.** Autophagy in plants: physiological roles and post-translational regulation. *Journal of Integrative Plant Biology* **63**: 161-179.

**Roberts MR, Salinas J, Collinge DB. 2002.** 14-3-3 proteins and the response to abiotic and biotic stress. *Plant Molecular Biology* **50**: 1031-1039.

**Saddhe AA, Mishra AK, Kumar K. 2021.** Molecular insights into the role of plant transporters in salt stress response. *Physiologia Plantarum* **173**: 1481-1494.

**Scharf KD, Berberich T, Ebersberger I, Nover L. 2012.** The plant heat stress transcription factor (Hsf) family: structure, function and evolution. *Biochimica Et Biophysica Acta-Gene Regulatory Mechanisms* **1819**: 104-119.

**Sebastian D, Fernando FD, Raul DG, Gabriela GM. 2020.** Overexpression of *Arabidopsis* aspartic protease *APA1* gene confers drought tolerance. *Plant Science* **292**: 110406.

**Shahinnia F, Geyer M, Schürmann F, Rudolphi S, Holzapfel J, Kempf H, Stadlmeier M, Löschenberger F, Morales L, Buerstmayr H, Sánchez JIy, Akdemir D, Mohler V, Lillemo M, Hartl L. 2022.** Genome-wide association study and genomic prediction of resistance to stripe rust in current Central and Northern European winter wheat germplasm. *Theoretical and Applied Genetics* **135**: 3583-3595.

**Sharma S, Prasad A, Sharma N, Prasad M. 2021.** Role of ubiquitination enzymes in abiotic environmental interactions with plants. *International Journal of Biological Macromolecules* **181**: 494-507.

**Sheng P, Tan J, Jin M, Wu F, Zhou K, Ma W, Heng Y, Wang J, Guo X, Zhang X, Cheng Z, Liu L, Wang C, Liu X, Wan J. 2014.** *Albino midrib 1*, encoding a putative potassium efflux antiporter, affects chloroplast development and drought tolerance in rice. *Plant Cell Reports* **33**: 1581-1594.

**Shi C, Baldwin IT, Wu JQ. 2012.** *Arabidopsis* plants having defects in nonsense-mediated mRNA decay factors UPF1, UPF2, and UPF3 show photoperiod-dependent phenotypes in development and stress responses. *Journal of Integrative Plant Biology* **54**: 99-114.

**Shin D, Koo YD, Lee J, Lee HJ, Baek D, Lee S, Cheon C, Kwak SS, Lee SY, Yun D. 2004.** Athb-12, a homeobox-leucine zipper domain protein from *Arabidopsis thaliana*, increases salt tolerance in yeast by regulating sodium exclusion. *Biochemical and Biophysical Research Communications* **323**: 534-540.

**Singh A, Pandey A, Srivastava AK, Tran LSP, Pandey GK. 2016.** Plant protein phosphatases 2C: from genomic diversity to functional multiplicity and importance in stress management. *Critical Reviews in Biotechnology* **36**: 1023-1035.

**Sohani MM, Schenk PM, Schultz CJ, Schmidt O. 2009.** Phylogenetic and transcriptional analysis of a strictosidine synthase-like gene family in *Arabidopsis thaliana* reveals involvement in plant defence responses. *Plant Biology* **11**: 105-117.

**Song ZT, Liu JX, Han JJ. 2021.** Chromatin remodeling factors regulate environmental stress responses in plants. *Journal of Integrative Plant Biology* **63**: 438-450.

**Stroher E, Millar AH. 2012.** The biological roles of glutaredoxins. *Biochemical Journal* **446**: 333-348.

**Su T, Li XZ, Yang MY, Shao Q, Zhao YX, Ma CL, Wang PP. 2020.** Autophagy: an intracellular degradation pathway regulating plant survival and stress response. *Frontiers in Plant Science* **11**: 784-91.

**Sun SH, Hu CG, Qi XJ, Chen JY, Zhong YP, Muhammad A, Lin MM, Fang JB. 2021.** The *AaCBF4-AaBAM3.1* module enhances freezing tolerance of kiwifruit (*Actinidia arguta*). *Horticulture Research* **8**: 97.

**Sun YY, Xu WZ, Wu L, Wang RZ, He ZY, Ma M. 2016.** An *Arabidopsis* mutant of inositol pentakisphosphate 2-kinase *AtIPK1* displays reduced arsenate tolerance. *Plant Cell and Environment* **39**: 416-426.

**Takahara K, Kasajima I, Takahashi H, Hashida S, Itami T, Onodera H, Toki S, Yanagisawa S, Kawai-Yamada M, Uchimiya H. 2010.** Metabolome and photochemical analysis of rice plants overexpressing *Arabidopsis* NAD kinase gene. *Plant Physiology* **152**: 1863-1873.

**Takahashi H, Watanabe A, Tanaka A, Hashida SN, Kawai-Yamada M, Sonoike K, Uchimiya H. 2006.** Chloroplast NAD kinase is essential for energy transduction through the xanthophyll cycle in photosynthesis. *Plant and Cell Physiology* **47**: 1678-1682.

**Trenner J, Monaghan J, Saeed B, Quint M, Shabek N, Trujillo M. 2022.** Evolution and functions of plant U-Box proteins: from protein quality control to signaling. *Annual Review of Plant Biology* **73**: 93-121.

**Unver T, Turktas M, Budak H. 2013.** In planta evidence for the involvement of a Ubiquitin Conjugating Enzyme (UBC E2 clade) in negative regulation of disease resistance. *Plant Molecular Biology Reporter* **31**: 323-334.

**Vessal S, Arefian M, Siddique KHM. 2020.** Proteomic responses to progressive dehydration stress in leaves of chickpea seedlings. *BMC Genomics* **21**: 523.

**Volz R, Kim SK, Mi JN, Mariappan KG, Guo XJ, Bigeard J, Alejandro S, Pflieger D, Rayapuram

**N, Al-Babili S, Hirt H. 2018.** The Trihelix transcription factor GT2-like 1 (GTL1) promotes salicylic acid metabolism, and regulates bacterial-triggered immunity. *PLoS genetics* **14**: e1007708.

**Wan JR, Tanaka K, Zhang XC, Son GH, Brechenmacher L, Tran HNN, Stacey G. 2012.** LYK4, a Lysin motif receptor-like kinase, is important for chitin signaling and plant innate immunity in *Arabidopsis*. *Plant Physiology* **160**: 396-406.

**Wan JR, Zhang XC, Neece D, Ramonell KM, Clough S, Kim SY, Stacey MG, Stacey G. 2008.** A LysM receptor-like kinase plays a critical role in chitin signaling and fungal resistance in *Arabidopsis*. *Plant Cell* **20**: 471-481.

**Wang H, Lu YQ, Jiang TT, Berg H, Li C, Xia YJ. 2013.** The *Arabidopsis* U-box/ARM repeat E3 ligase AtPUB4 influences growth and degeneration of tapetal cells, and its mutation leads to conditional male sterility. *Plant Journal* **74**: 511-523.

**Wang H, Xu Q, Kong YH, Chen Y, Duan JY, Wu WH, Chen YF. 2014.** Arabidopsis WRKY45 transcription factor activates *PHOSPHATE TRANSPORTER1;1* expression in response to phosphate starvation. *Plant Physiology* **164**: 2020-2029.

**Wang QQ, Liu JY, Wang Y, Zhao Y, Jiang HY, Cheng BJ. 2015.** Systematic analysis of the maize PHD-finger gene family reveals a subfamily involved in abiotic stress response. *International Journal of Molecular Sciences* **16**: 23517-23544.

**Wang R, Fang YN, Wu XM, Qing M, Li CC, Xie KD, Deng XX, Guo WW. 2020.** The miR399-CsUBC24 module regulates reproductive development and male fertility in *Citrus*. *Plant Physiology* **183**: 1681-1695.

**Wang S, Lv XY, Zhang JL, Chen DN, Chen SX, Fan GQ, Ma CQ, Wang YG. 2022.** Roles of E3 Ubiquitin ligases in plant responses to abiotic stresses. *International Journal of Molecular Sciences* **23**: 2308.

**Wang XM, Shi X, Chen SY, Ma C, Xu SB. 2018.** Evolutionary origin, gradual accumulation and functional divergence of Heat Shock Factor gene family with plant evolution. *Frontiers in Plant Science* **9**: 71.

**Wang Y, Cai SY, Yin LL, Shi K, Xia XJ, Zhou YH, Yu JQ, Zhou J. 2015.** Tomato HsfA1a plays a critical role in plant drought tolerance by activating *ATG* genes and inducing autophagy. *Autophagy* **11**: 2033-2047.

**Wei M, Zhuang Y, Li H, Li PH, Huo HQ, Shu D, Huang WZ, Wang SH. 2020.** The cloning and characterization of *hypersensitive to salt stress* mutant, affected in quinolinate synthase, highlights the involvement of NAD in stress-induced accumulation of ABA and proline. *Plant Journal* **102**: 85-98.

**Xing S, Zachgo S. 2008.** *ROXY1* and *ROXY2*, two *Arabidopsis* glutaredoxin genes, are required for anther development. *Plant Journal* **53**: 790-801.

**Xiong Y, Contento AL, Bassham DC. 2005.** AtATG18a is required for the formation of autophagosomes during nutrient stress and senescence in *Arabidopsis thaliana*. *Plant Journal* **42**: 535-546.

**Xu J, Zhang SQ. 2015.** Mitogen-activated protein kinase cascades in signaling plant growth and development. *Trends in Plant Science* **20**: 56-64.

**Xu WF, Shi WM, Jia LG, Liang JS, Zhang JH. 2012.** TFT6 and TFT7, two different members of tomato 14-3-3 gene family, play distinct roles in plant adaption to low phosphorus stress. *Plant Cell and Environment* **35**: 1393-1406.

**Xu XM, Zhang LL, Zhao W, Fu L, Han YX, Wang KK, Yan LY, Li Y, Zhang XH, Min DH. 2021.** Genome-wide analysis of the serine carboxypeptidase-like protein family in *Triticum aestivum* reveals TaSCPL184-6D is involved in abiotic stress response. *BMC Genomics* **22**: 350.

**Yamaguchi YL, Ishida T, Sawa S. 2016.** CLE peptides and their signaling pathways in plant development. *Journal of Experimental Botany* **67**: 4813-4826.

**Yang YW, Yang LT, Li ZG. 2013.** Molecular cloning and identification of a putative tomato cationic amino acid transporter-2 gene that is highly expressed in stamens. *Plant Cell Tissue and Organ Culture* **112**: 55-63.

**Yao X, Xiong W, Ye TT, Wu Y. 2012.** Overexpression of the aspartic protease *ASPG1* gene confers drought avoidance in *Arabidopsis*. *Journal of Experimental Botany* **63**: 2579-2593.

**Yao Y, Zhou JH, Cheng C, Niu FA, Zhang AP, Sun B, Tu RJ, Wan JN, Li Y, Huang YW, Xie KZ, Dai YT, Zhang H, Hong JH, Pan XH, Zhu JJ, Zhou H, Liu ZH, Cao LM, Chu HW. 2022.** A conserved clathrin-coated vesicle component, OsSCYL2, regulates plant innate immunity in rice. *Plant Cell and Environment* **45**: 542-555.

**Ye Y, Ding Y, Jiang Q, Wang F, Sun J, Zhu C. 2017.** The role of receptor-like protein kinases (*RLKs*) in abiotic stress response in plants. *Plant Cell Reports* **36**: 235-242.

**Yuenyong W, Sirikantaramas S, Qu LJ, Buaboocha T. 2019.** Isocitrate lyase plays important roles in plant salt tolerance. *BMC Plant Biol* **19**: 472.

**Zhang J, Feng JJ, Lu J, Yang YZ, Zhang X, Wan DS, Liu JQ. 2014.** Transcriptome differences between two sister desert poplar species under salt stress. *BMC Genomics* **15**: 337.

**Zhang MM, Zhang SQ. 2022.** Mitogen-activated protein kinase cascades in plant signaling. *Journal of Integrative Plant Biology* **64**: 301-341.

**Zhang TE, Li XM, Zhao Q, Shi Y, Hao YJ, You CX. 2022.** Genome-wide identification and functional characterization of the MdCLE peptide family in apple ( *Malus x domestica* ). *Horticultural Plant Journal* **8**: 279-288.

**Zhang WD, Wang L, Wang YZ, Wang Y, Gao QR. 2020.** Cloning and characterization of *TaPUB4*, a U-box gene from common wheat (*Triticum aestivum* L.) involved in regulation of pollen development by influencing sucrose-starch pathway in anther. *Molecular Breeding* **40**: 56.

**Zhang YR, Zhou JF, Wei F, Song TQ, Yu Y, Yu M, Fan QR, Yang YN, Xue G, Zhang XK. 2021.** Nucleoredoxin gene *TaNRX1* positively regulates drought tolerance in transgenic Wheat (*Triticum aestivum* L.). *Frontiers in Plant Science* **12**: 756338.

**Zhang YW, Li ZW, Ma B, Hou QC, Wan XY. 2020.** Phylogeny and Functions of LOB domain proteins in plants. *International Journal of Molecular Sciences* **21**: 2278.

**Zhang ZB, Hu MH, Feng XB, Gong AD, Cheng L, Yuan HY. 2017.** Proteomes and phosphoproteomes of anther and pollen: availability and progress. *Proteomics* **17**: 20.

**Zhao MW, Ge Y, Xu ZY, Ouyang X, Jia YL, Liu JT, Zhang MX, An YY. 2022.** A BTB/POZ domain-containing protein negatively regulates plant immunity in *Nicotiana benthamiana*. *Biochemical and Biophysical Research Communications* **600**: 54-59.

**Zhou XM, Zhao P, Wang W, Zou J, Cheng TH, Peng XB, Sun MX. 2015.** A comprehensive, genome-wide analysis of autophagy-related genes identified in tobacco suggests a central role of autophagy in plant response to various environmental cues. *DNA Research* **22**: 245-257.

**Zhu ML, Bin J, Ding HF, Pan D, Tian QY, Yang XL, Wang LG, Yue YZ. 2022.** Insights into the trihelix transcription factor responses to salt and other stresses in *Osmanthus fragrans*. *BMC Genomics* **23**: 334.

**Zhu XJ, Pan T, Zhang X, Fan LG, Quintero FJ, Zhao H, Su XM, Li XJ, Villalta I, Mendoza I, Shen JB, Jiang LW, Pardo JM, Qiu QS. 2018.** $K^+$ efflux antiporters 4, 5, and 6 mediate pH and $K^+$ homeostasis in endomembrane compartments. *Plant Physiology* **178**: 1657-1678.
