## Supplementary Table S13 for "Positive selection and relaxed purifying selection contribute to rapid evolution of male-biased genes in a dioecious flowering plant"

**Table S13** Functions and references associated with abiotic stress and immune responses, organ developments of male-biased genes under significant relaxed selection (*P-value* < 0.05) in floral buds.

| OGs | Function Description | Signal* | Stress responses/Developments | References |
|---|---|---|---|---|
| OG0008899 | Non-canonical poly(A) RNA polymerase protein Trf4 | R | | |
| **OG0007568** | quinolinate synthase (QS) | R | salt | Wei *et al.* (2020) |
| **OG0008786** | basic 7S globulin-like protein | R | antibacterial, allergen, heat | Hirano (2021) |
| **OG0008247** | Homogentisate phytyltransferase 1 (HPT1) | R | light, oxidative, nutrient starvation | Collakova and DellaPenna (2003) |
| **OG0006131** | LOB domain-containing protein 18 (LBD18) | R | heat, light, pathogens, lateral roots formation, callus formation | Zhang *et al.* (2020); Liu *et al.* (2019) |
| **OG0007549** | WRKY transcription factor 72 (WRKY72) | R | salt, drought, heat, ABA, osmotic, pathogens, sugar starvation | Chen *et al.* (2017); Chen *et al.* (2012); Li *et al.* (2020) |
| ***OG0008560*** | CLAVATA3/ESR (CLE)-related protein 13(CLV3/CLE13) | R | cold, heat, drought, salt, cytokinin, hormonal, floral organ, stamen, anther, root, shoot | Zhang *et al.* (2022); Lin *et al.* (2021); Cui *et al.* (2022); Jun *et al.* (2010); Laffont *et al.* (2020) |
| **OG0009366** | glycosyltransferase family protein 64 C3 (GT64-C3) | R | terrestrial environment | Caputi *et al.* (2012) |
| **OG0008638** | E3 ubiquitin-protein ligase AIRP2 (AIRP2) | R | abscisic acid (ABA), drought | Yu *et al.* (2016); Cho *et al.* (2011) |
| OG0008572 | uncharacterized protein | R | | |
| **OG0008611** | AUXIN SIGNALING F-BOX 2 (AFB2) | R | drought, salt, oxidative | Verma *et al.* (2022); Iglesias *et al.* (2010) |
| **OG0007091** | ethylene-responsive transcription factor ERF060 | R | alpine environment | Ma *et al.* (2015) |
| **OG0008547** | phosphorelay response regulator | R | cytokinins, ethylene, abscisic | Skalak *et al.* |

| | | | | |
|---|---|---|---|---|
| | | | acid, light, temperature | (2021) |
| OG0006106 | uncharacterized protein | R | | |
| **OG0006323** | Glycosyltransferase family 92 protein (GT92) | R | terrestrial environment | Ebert *et al.* (2018); Caputi *et al.* (2012) |
| **OG0007242** | subtilisin-like protease SBT1.4 | R | osmotic | Stührwohldt *et al.* (2021) |
| OG0009543 | mediator of RNA polymerase II transcription subunit 25 | R | | |
| ***OG0008341*** | pollen receptor-like kinase 3 (PRK3) | R | pollen development, pollen grain, pollen maturation | Kim *et al.* (2021); Muschietti and Wengier (2018); Takeuchi and Higashiyama (2016); Kim *et al.* (2002) |

OGs in bold font are associated with biotic and abiotic stress responses. OGs in bold and italic fonts are associated with pollen developments and maturation.

Asterisk (*) indicates that relaxed selections are tested by RELAX (R).


**Caputi L, Malnoy M, Goremykin V, Nikiforova S, Martens S. 2012.** A genome-wide phylogenetic reconstruction of family 1 UDP-glycosyltransferases revealed the expansion of the family during the adaptation of plants to life on land. *Plant Journal* **69**: 1030-1042.

**Chen F, Hu Y, Vannozzi A, Wu KC, Cai HY, Qin Y, Mullis A, Lin ZG, Zhang LS. 2017.** The WRKY transcription factor family in model plants and crops. *Critical Reviews in Plant Sciences* **36**: 311-335.

**Chen LG, Song Y, Li SJ, Zhang LP, Zou CS, Yu DQ. 2012.** The role of WRKY transcription factors in plant abiotic stresses. *Biochimica Et Biophysica Acta-Gene Regulatory Mechanisms* **1819**: 120-128.

**Cho SK, Ryu MY, Seo DH, Kang BG, Kim WT. 2011.** The Arabidopsis RING E3 Ubiquitin Ligase AtAIRP2 plays combinatory roles with AtAIRP1 in abscisic acid-mediated drought stress responses. *Plant Physiology* **157**: 2240-2257.

**Collakova E, DellaPenna D. 2003.** The role of homogentisate phytyltransferase and other tocopherol pathway enzymes in the regulation of tocopherol synthesis during abiotic stress. *Plant Physiology* **133**: 930-940.

**Cui YW, Lu XT, Gou XP. 2022.** Receptor-like protein kinases in plant reproduction: current understanding and future perspectives. *Plant Communications* **3:** 100273.

**Ebert B, Birdseye D, Liwanag AJM, Laursen T, Rennie EA, Guo XY, Catena M, Rautengarten C, Stonebloom SH, Gluza P, Pidatala VR, Andersen MCF, Cheetamun R, Mortimer JC, Heazlewood JL, Bacic A, Clausen MH, Willats WGT, Scheller HV. 2018.** The three members of the Arabidopsis glycosyltransferase family 92 are functional beta-1,4-Galactan synthases. *Plant and Cell Physiology* **59**: 2624-2636.

**Hirano H. 2021.** Basic 7S globulin in plants. *Journal of Proteomics* **240**: 104209.

**Iglesias MJ, Terrile MC, Bartoli CG, DIppolito S, Casalongue CA. 2010.** Auxin signaling participates in the adaptative response against oxidative stress and salinity by interacting with redox metabolism in *Arabidopsis*. *Plant Molecular Biology* **74**: 215-222.

**Jun J, Fiume E, Roeder AHK, Meng L, Sharma VK, Osmont KS, Baker C, Ha CM, Meyerowitz EM, Feldman LJ, Fletcher JC. 2010.** Comprehensive analysis of CLE polypeptide signaling gene expression and overexpression activity in *Arabidopsis*. *Plant Physiology* **154**: 1721-1736.

**Kim HU, Cotter R, Johnson S, Senda M, Dodds P, Kulikauskas R, Tang WH, Ezcurra I, Herzmark P, McCormick S. 2002.** New pollen-specific receptor kinases identified in tomato, maize and Arabidopsis: the tomato kinases show overlapping but distinct localization patterns on pollen tubes. *Plant Molecular Biology* **50**: 1-16.

**Kim MJ, Jeon BW, Oh E, Seo PJ, Kim J. 2021.** Peptide signaling during plant reproduction. *Trends in Plant Science* **26**: 822-835.

**Laffont C, Ivanovici A, Gautrat P, Brault M, Djordjevic MA, Frugier F. 2020.** The NIN transcription factor coordinates CEP and CLE signaling peptides that regulate nodulation antagonistically. *Nature Communication* **11**: 3167.

**Li WX, Pang SY, Lu ZG, Jin B. 2020.** Function and mechanism of WRKY transcription factors in abiotic stress responses of plants. *Plants* **9**: 1515.

**Lin H, Wang W, Chen XG, Sun ZT, Han XL, Wang S, Li Y, Ye WW, Yin ZJ. 2021.** Molecular traits and functional analysis of the CLAVATA3/Endosperm surrounding region-related small signaling peptides in three species of *Gossypium* Genus. *Frontiers in Plant Science* **12**: 671626.

**Liu SQ, Wang B, Li XJ, Pan JX, Qian XX, Yu YH, Xu P, Zhu J, Xu XF. 2019.** Lateral Organ Boundaries Domain 19 (LBD19) negative regulate callus formation in *Arabidopsis*. *Plant Cell Tissue and Organ Culture* **137**: 485-494.

**Ma L, Sun XD, Kong XX, Galvan JV, Li X, Yang SH, Yang YQ, Yang YP, Hu XY. 2015.** Physiological, biochemical and proteomics analysis reveals the adaptation strategies of the alpine plant *Potentilla saundersiana* at altitude gradient of the Northwestern Tibetan Plateau. *Journal of Proteomics* **112**: 63-82.

**Muschietti JP, Wengier DL. 2018.** How many receptor-like kinases are required to operate a pollen tube. *Current Opinion in Plant Biology* **41**: 73-82.

**Skalak J, Nicolas KL, Vankova R, Hejatko J. 2021.** Signal integration in plant abiotic stress responses via multistep phosphorelay signaling. *Frontiers in Plant Science* **12**: 644823.

**Stührwohldt N, Bühler E, Sauter M, Schaller A. 2021.** Phytosulfokine (PSK) precursor processing by subtilase SBT3.8 and PSK signaling improve drought stress tolerance in *Arabidopsis*. *Journal of experimental botany* **72**: 3427-3440.

**Takeuchi H, Higashiyama T. 2016.** Tip-localized receptors control pollen tube growth and LURE sensing in *Arabidopsis*. *Nature* **531**: 245-8.

**Verma S, Negi NP, Pareek S, Mudgal G, Kumar D. 2022.** Auxin response factors in plant adaptation to drought and salinity stress. *Physiologia Plantarum* **174**: e13714.

**Wei M, Zhuang Y, Li H, Li PH, Huo HQ, Shu D, Huang WZ, Wang SH. 2020.** The cloning and characterization of hypersensitive to salt stress mutant, affected in quinolinate synthase, highlights the involvement of NAD in stress-induced accumulation of ABA and proline. *Plant Journal* **102**: 85-98.

**Yu FF, Wu YR, Xie Q. 2016.** Ubiquitin-proteasome system in ABA signaling: from perception to action. *Molecular Plant* **9**: 21-33.

**Zhang TE, Li XM, Zhao Q, Shi Y, Hao YJ, You CX. 2022.** Genome-wide identification and functional characterization of the MdCLE peptide family in apple (*Malus x domestica*). *Horticultural Plant Journal* **8**: 279-288.

**Zhang YW, Li ZW, Ma B, Hou QC, Wan XY. 2020.** Phylogeny and functions of LOB domain proteins in plants. *International Journal of Molecular Sciences* **21**: 2278.
